# Museums and Zoos: Rapid genetic identification of rare species and practical applications for conservation and systematics in a biodiverse country

**DOI:** 10.1101/2025.02.10.637512

**Authors:** Daniel E. Chavez, Julio C. Carrión-Olmedo, María B. Cabezas, Daniela Reyes-Barriga, Pamela Lojan, David Mora, Martin Bustamante, C. Miguel Pinto, Pablo Jarrin-V

## Abstract

Obtaining genetic information from rare species is challenging for scientists, but it is crucial for understanding animal evolutionary history and informing conservation management initiatives. We present the first example of a collaborative local network that includes zoos and natural history collections to investigate the evolution, systematics, and conservation concerns of olingos (genus *Bassaricyon*, Procyonidae, Carnivora, Mammalia). We sequenced the entire (1,146 base pairs) cytochrome *b* gene to phylogenetically identify individuals that have been victims of wildlife trafficking. Unexpectedly, we detected an individual specimen belonging to *Bassaricyon medius orinomus* (western lowland olingo), which may represent a new geographical record for this taxon in Ecuador. Through our practical experiences, we describe how local collaboration is possible and crucial for promoting wildlife genetic research in the Global South and contributing to protecting the last populations of rare mammals. We also discuss the significance of wild animals under human care as a valuable genetic resource for scientific research, conservation strategies, and informed wildlife management decisions.

## Introduction

Obtaining biological material from uncommon and endangered species is a challenging scientific endeavor, yet, the availability of such material is crucial for successful conservation efforts (Consortium, 2020; Ekblom et al., 2018; Khan et al., 2016; Rubinoff et al., 2020). As a “repository and living record of the world’s biodiversity”, zoos are important national resources for research on rare species, for supporting wildlife conservation, and as a source of special holdings in natural history collections (Poo et al., 2022; Powell et al., 2023). With the progress of science and technology, zoos have transformed into essential research hubs, contributing significantly to biotechnology and biomedicine (Andrews et al., 2023; Consortium, 2020; Powell et al., 2023; Sullivan et al., 2023). Research from zoo animals has contributed to understanding the impact of climate change, the potential of anti-microbial proteins, the mechanisms behind human disease, and how cancerous tumors can be suppressed (Powell et al., 2023; Sullivan et al., 2023). By studying animal physiology and biomolecules, zoos play a crucial role in advancing scientific knowledge, improving animal care, and contributing to conservation efforts (Consortium, 2020; Patrick & Tunnicliffe, 2013). Particularly, zoos house a substantial portion of threatened species including 9% of reptiles, 42% of mammals, 23% of birds, 9% of amphibians, and 7% of fish (Species360, 2024).

Despite the importance of genetic research on rare species, one significant challenge is exporting biological material for DNA sequencing across national frontiers (Alves et al., 2018). Notably, international and local government regulations that require permits to export biological material from the country of origin can take long periods to acquire (Lawson et al., 2022; Vilaça et al., 2024). Additionally, conducting genetic analyses abroad can lead to project delays and compromised sample quality due to unexpected losses or degradation of samples along the transport route (Vilaça et al., 2024). Outsourcing sample analysis beyond the national frontiers of the source, usually a developing and biodiverse country, generates additional problems related to scientific development, knowledge production, and technological dependency (Brichieri-Colombi et al., 2019; Díaz et al., 2021; Jarrín-V et al., 2021).

Newer and cheaper technologies have facilitated local initiatives for nucleic acid sequencing in the Global South (Pomerantz et al., 2018; Wang et al., 2021). High-throughput sequencing has become popular in developing countries, creating opportunities to improve genetic research (Jayathissa & Rupasinghe, 2024; Pozo et al., 2024). However, the contribution to species conservation through access to biological material from zoos remains wanting in developing countries. This is related to the lack of necessary financial resources to properly store biological samples and provide DNA or next-generation sequencing services and the availability of local expertise to analyze genomic data (Díaz et al., 2021; Jarrín-V et al., 2021). One solution for conservation genetics in the “Global South” (Dados & Connell, 2012), where resources are scarce and investment risk is high (Bolaños-Villegas et al., 2020), is to create a decentralized Biobank consortium that includes zoos, natural history collections, specialized laboratories, and universities (Dumancas et al., 2023). Decentralization should help to include more stakeholders, adding expertise from multiple institutions and increasing chances for funding and community engagement (Weidener & Spreckelsen, 2024).

In the essential application of the identification of individuals and species that are the victims of illegal wildlife trafficking, the genetic material from zoos provides a unique opportunity to address two concerns (Consortium, 2020; Miranda et al., 2015; Miranda et al., 2023; Powell et al., 2023). (1) DNA can determine the identity of cryptic species and, in some cases, allow inference of geographic origin for a given specimen. This helps identify the source population for species reintroduction and, in some cases, it may help identify poaching hotspots and understand illegal trade routes (Pérez-Espona & Consortium, 2021). (2) Adding new biological material from zoos into biobanks and natural history collections provides insights into the evolutionary history of species (Shaw et al., 2024).

Our study involved four individuals of olingos (genus *Bassaricyon*, Procyonidae) rescued from illegal mascotism (a form of wildlife trafficking in which local inhabitants take wild animals from their surroundings to keep them as pets). These four animals have been maintained in the Fundación Zoológica de Quito (Quito Zoo) since 2018. Because of their uncertain origin and young ages, morphological identifications were ambiguous. We present insights into the phylogeny of olingos (*Bassaricyon*), a group of charismatic carnivores that suffer conservation concerns. Here, we demonstrate how scientific networking among live collections at Quito Zoo and museum collections from the Instituto Nacional de Biodiversidad (INABIO) was crucial, not just for rapid molecular species identification, but also for informing the long-term conservation of *Bassaricyon* species in Ecuador.

Olingos represent an ideal group to test the benefits of a local collaboration network. On the one hand, olingos are relatively rare in captivity (Species360, 2024), making them coveted objects for the pet trade and also occasionally trafficked due to confusion with kinkajous (*Potos flavus*), which are more widely recognized and sought after in the exotic pet trade (Bush et al., 2014; Harrington, 2015). Investigating the geographic origin of the rescued olingos maintained at the Quito Zoo could provide insights into wildlife trafficking routes and opportunities for species reintroduction. On the other hand, olingos are uncommon in Latin American natural history collections. Only 13 specimens are present in museum collections in Latin America, representing only 15% of the collection worldwide (GBIF.org, 2025; Helgen et al., 2013). The low number of olingo specimens in the museum collections of the Global South limits research opportunities for understanding the biology and ecological significance of this group of mammals.

## Results

To determine the species identity of the olingo individuals housed at the Quito Zoo, we inferred a phylogeny based on complete sequences of the mitochondrial cytochrome *b* gene (*CYTB*; 1,146 bp) that included the four individuals that were hosted at the zoo. We compared these sequences with those from three additional samples deposited in the mammal collection of the INABIO, and 17 samples available in GenBank. To root the phylogenetic tree, we included sequences from *Nasua nasua*, *Procyon lotor*, and *Bassariscus sumichrasti* as outgroups. Collectively, the zoo and museum samples were distributed throughout the olingo clade of the inferred phylogeny, with the exception of the *B. gabbii* lineage, which is distributed from Nicaragua to western was Colombia only (Saavedra-Rodríguez & Velandia-Perilla, 2011).

Four samples belonged to the *B. medius* clade (Figure 1 and Figure S1). Interestingly, one sample from the Quito Zoo (named by the zoo as “Munay”) belonged to the *B. medius orinomus* subclade (Figure 1 and Figure 2), which is a subspecies assumed to be restricted to Panama and has never been recorded in Ecuador. The mean divergence time of this subclade was estimated to be 1.8 million years ago (mya) with a 95% highest posterior density (HPD) 0.9-2.6 million mya (Figure S1 and Table S1). The remaining three samples corresponding to sequences of the other three olingos at the Quito Zoo (“Clarissa”, “Macho”, and “Lola”) were positioned in a monophyletic clade with sequences representing the *B. medius medius* subspecies from the Choco region in Ecuador (Figure 1). Together, the *B. medius* sequences diverged from the *B. alleni* clade at 2.3 mya (95% HPD = 1.7 - 2.9 mya). The *B. alleni* clade included one sequence from the INABIO museum, MECN8080, and that clade showed a basal split at 1.7 mya (95% HPD = 0.4-2.8 mya). Finally, we recovered only one sequence from GenBank belonging to *B. gabbii*, with this diverging from the *B. alleni* + *B. medius* clade about 4 million years ago (95% HPD = 2.5 - 6.2 mya). The deepest clade within the olingo group comprises *B. neblina*, with an ancient divergence of 5.6 mya (95% HPD = 3.2 - 8.1 mya). This clade included one sequence, “TACU”, from the INABIO collection (Figure 1 and Figure S1).

**Figure 1.**
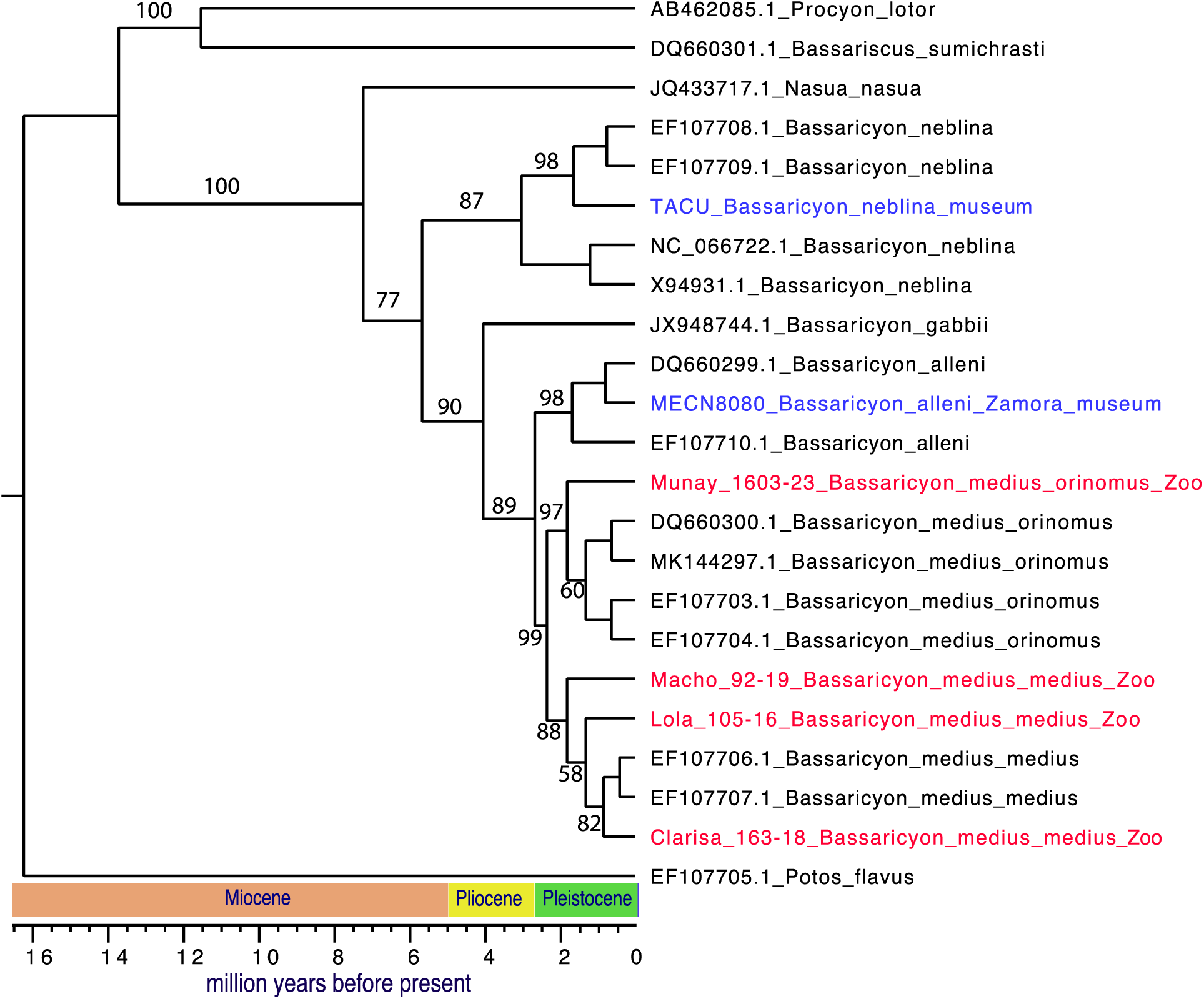
Inferred maximum likelihood tree and chronogram which includes the studied samples of olingos. See Figure S1 and Table S1 for estimates with 95% highest posterior density intervals and Table S2 for fossil priors. A total of 23 sequences were included in the analysis, with four individuals from the Quito Zoo and two individuals from the INABIO natural history collection. Blue-labeled branches are samples from INABIO. Red-labeled branches are sequences from individuals under human care at the Quito Zoo. Bootstrap support values (out of 1000 ultrafast bootstrap replicates) are shown at the tree nodes.

**Figure 2.**
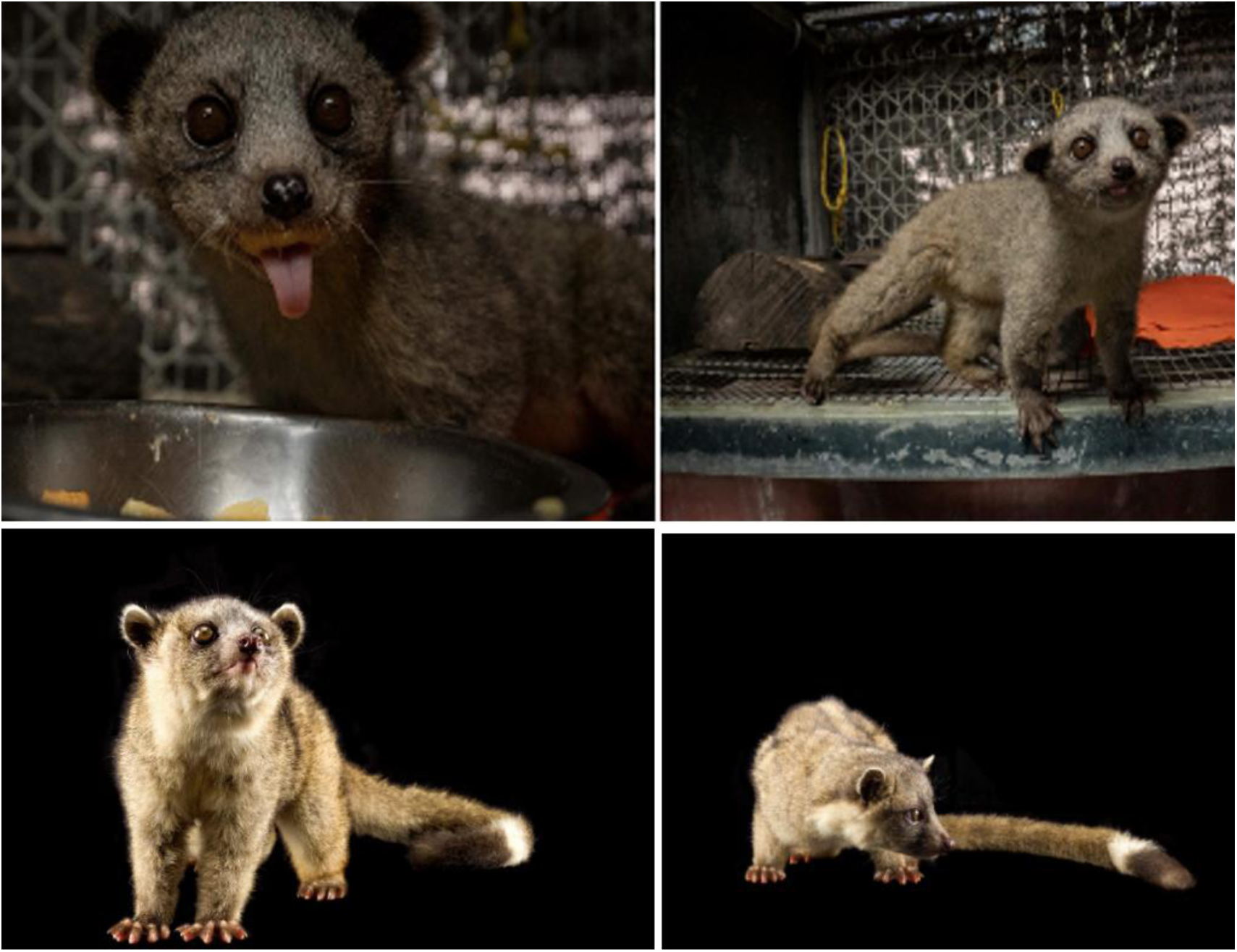
Top row: photos of “Munay” (identified as belonging to *Bassaricyon medius orninomus*). This animal represents either a new geographic record or a victim of international wildlife trafficking. Bottom row: photos of “Clarissa” (identified as belonging to *Bassaricyon medius medius*). This species has been previously recorded for Ecuador. Photos were taken by Andy Reinoso © 2021 and used in this work with permission from the author.

## Discussion

### Evolutionary history of olingos in South America

The inclusion of samples from both zoos and natural history collections allowed us to investigate the evolution of the genus *Bassaricyon* in South America. Our estimates showed that the ancestor of olingos existed around 7.2 mya (Figure 1 and Table S1). This is consistent with a dispersal event of this group through island hopping (Helgen et al., 2013; Koepfli et al., 2007). In particular, at that time, the Panamanian land bridge was not fully formed, and this region was a vast archipelago (Webb, 1991; Webb, 2006). During this time, the ancestor species of olingos may have dispersed through different islands when sea levels were relatively low (Coates & Obando, 1996; O’Dea et al., 2016; Webb & Rancy, 1996). Concordant with immigration into South America via island hopping, our phylogenetic tree showed that the earliest divergence within the genus was between *B. neblina* and the remaining extant *Bassaricyon* species around 6 mya (Figure 1 and Table S1), which is consistent with previous estimates (Helgen et al., 2013).

*B. neblina* is distributed in the central montane region of the Andes, which suggests that the ancestor of olingos may have used a natural corridor known as the “Andean route” to disperse from Central America across the Andes (Berta, 1987; Webb, 1978). The route was followed by other carnivores (Berta, 1987; Chavez et al., 2022; Eizirik et al., 2010). Soon after, around 4 mya, olingos may have dispersed from the central region of the Andes to the eastern slopes to give rise to *B. gabbii* (Helgen et al., 2013). Then, at around 2.6 mya, olingos most likely reached the lowlands of South America to give rise to *B. medius* in the western region and *B. alleni* in the eastern part of the Andes (Figure 1 and Table S1). Our results showed rapid speciation of olingos during the Pleistocene, most likely due to available niches in South America (Koepfli et al., 2007) and the Andes acting as a barrier to gene flow and contributing to diversification (Helgen et al., 2013).

### New geographical records and illegal trade routes

The integration of rare and endangered species samples from zoos into natural history collections provides a unique opportunity for advancing knowledge, including new geographical records (Poo et al., 2022; Powell et al., 2023). We have illustrated this by analyzing the records of *Bassaricyon medius* in the Quito Zoo. This species is distributed in the lower slopes of the western side of the Andes of Ecuador and Colombia, and north to Panama (Chacón-Pacheco et al., 2019; Helgen, 2016; Ramírez-Chaves et al., 2022). Two subspecies have been described that have been suggested to have little geographic overlap: *B. medius medius* in Ecuador and Colombia, and *B. medius orinomus* in Panama and potentially northern Colombia (Helgen et al., 2013). However, the record of a *B. medius orinomus* from the Quito Zoo (“Munay” in Figure 1 and Figure 2) challenges our knowledge of the distribution of this group if indeed this animal originated from Ecuador. Alternatively, this animal could have been trafficked from Panamá or northern Colombia.

The animal in question, a young male, was rescued from a household in the La Esperanza Community, near the town of Rio Verde, in the parish of Lita, in the Imbabura Province, an underserved settlement in northwestern Ecuador. Thus, it is most likely that this olingo was a victim of local animal trafficking rather than being brought across international borders. Therefore, if *B. medius orinomus* is naturally present in Ecuador, it may largely overlap with *B. medius medius* along the latter species’ distribution (Chacón-Pacheco et al., 2019; Helgen et al., 2013; Ramírez-Chaves et al., 2022). Both subspecies are reciprocally monophyletic (Figure 1); yet the possibility of interbreeding cannot be rejected, as our analyses are based on a single gene of the mitochondrial genome. We recommend more genetic surveys, particularly using nuclear markers, to assess the integrity of both subspecies. We also know that, on average, *B. medius orinomus* is larger than *B. medius medius* (Helgen et al., 2013); thus, ecological studies are required to understand how these species interact in overlapping parts of their geographic distributions.

### Collaboration between zoos and natural history collections is critical for the rapid sequencing of rare species

Our current work has demonstrated that animals under the guardianship of zoos can provide significant genetic resources for evolutionary history and conservation genetics studies (Consortium, 2020; Poo et al., 2022; Powell et al., 2023), particularly in biodiverse and poorly studied regions of the world. Often, these animals arrive at zoos as victims of trafficking and, therefore, belong to the part of biodiversity that is under threat. These animals also contribute to strengthening information from the specimens available in museum collections (Poo et al., 2022; Powell et al., 2023). Notably, many animals in zoos represent species that are rare, endangered, or even extinct in the wild (Che-Castaldo et al., 2018). Collecting samples from these specimens in the wild could be difficult and expensive (Powell et al., 2023). For instance, one of the authors and his research team required several months and about $25,000 USD (Pinto, com. pers.) to find one live specimen in the olingo group, an effort that later resulted in novel insights on hidden biodiversity (Helgen et al., 2013). In contrast to this previous experience, we obtained new biological material and insights at no cost, as the sources were collected during routine health assessments at the Quito Zoo. Recognizing that the DNA of rare species is difficult to obtain in the field, that most scientists do not have the financial resources to cover expensive and prolonged field expeditions, and the urgent need to understand and protect species under extinction threat, zoos must conserve as much of the biological samples from their species as possible (Mooney et al., 2023). However, the proper storage of blood and tissues requires applying the principles of a biobank, including considerable investment and planning, data management strategies, freezer space, and trained staff (Poo et al., 2022; Powell et al., 2023). Collaboration becomes essential in the Global South, where zoos and museum collections struggle for adequate funding. Therefore, our work demonstrates a successful exercise where museums and zoos can supplement the requirements of the other.

The usefulness of DNA from zoos has been extensively demonstrated (Andrews et al., 2023; Christmas et al., 2023; Foley et al., 2023; Sullivan et al., 2000). Specifically, the availability of frozen biological material has allowed a detailed understanding of mammalian genomes and their implications in evolution, human diseases, and conservation through the Zoonomia Consortium (Christmas et al., 2023; Sullivan et al., 2000; Wilder et al., 2023). Since the genome assembly data generated by the Zoonomia Consortium included many individuals born under human care in zoos from the Global North, the application of such a database is limited concerning the evaluation of inbreeding and genetic diversity in the wild (Poo et al., 2022). In contrast, in the Global South, zoos and rescue centers are valuable resources for conservation genetics as most individuals come from the wild. Our olingo samples contain yet-to-be-analyzed whole genomes to infer demographic history, genome-wide genetic diversity, and quantify the amounts of deleterious variation (Chavez et al., 2022; Robinson et al., 2023). This information could assist researchers in evaluating the risk of extinction of olingos and optimizing and informing conservation management strategies like genetic rescue (Kyriazis et al., 2023; Robinson et al., 2022).

Despite the importance of incorporating rare species into natural history collections, some museums may consider zoo specimens of low scientific value because of the lack of specific geographical origin data and coordinates. Moreover, samples obtained from traffickers includes the condition of forced anonymity and lack of source data (Powell et al., 2023). We recognize that this information gap may negatively impact some aspects of the scientific value of zoo specimens. However, there is considerable value for demography and translocation programs if geographic information is known to some extent (He et al., 2016; Marr et al., 2018; Powell et al., 2023) or if it could be inferred through phylogeographic and population genomic analysis. For instance, we identified two subspecies of *B. medius* among individuals from the Quito Zoo (Figure 1). Individuals belonging to *B. medius medius* are distributed in the Choco region in Northern Ecuador (Chacón-Pacheco et al., 2019; Ramírez-Chaves et al., 2022) and could be potentially reintroduced in that region. In contrast, the other subspecies *B. medius orinomus* has only been reported in the Panama region (Helgen, 2016; Ramírez-Chaves et al., 2022), so further research is required before reintroducing this individual in Ecuador.

Another major challenge that Ecuadorian zoos face in determining species and possible origins via genetic data is the time and cost of DNA sequencing. Traditionally, DNA sequencing in Ecuador has been conducted outside the country (for example see Ortega et al., 2025). However, with recently acquired technologies for local sequencing in Ecuadorian natural history collections (Brito et al., 2024), we have found that the price to sequence the *CYTB* gene from one sample is 12 times less than sending samples to international companies (pers. observation). In particular, the cost to sequence an average gene of 1kb in length with Sanger sequencing abroad is ∼$20 USD ($10 USD forward plus $10 USD reverse). In contrast, the cost to sequence a gene with an Oxford Nanopore Technologies instrument locally is only ∼$1.73 USD. This estimate is based on adding $0.70 for sequencing (estimated cost of a Flongle flow cell per barcode) plus $0.78 for library prep (cost-kit/ per barcode) and $0.25 cents for expendables (e.g., laboratory plastic tubes), and because the sequencing technicians are part of the research team, labor costs are subsidized as part of the collaboration. On the other hand, shipping samples to other countries generates at least a 30-day increase in the waiting time to receive the data and adds courier costs and risks (e.g., sample degradation or loss during shipment). Furthermore, the required export time can be extended up to at least 50 days or more if a CITES permit is required by the importing country. Overall, we strongly believe that generating data with available technologies within a country such as Ecuador, despite the challenges, often has an intangible value for local development and opportunity (Jarrín-V et al., 2021).

### The construction of a local biobank for managing zoo genetic resources and species reintroduction

In this study, we have identified four distinct lineages of olingos from wild populations in Ecuador based on samples either collected from zoo animals confiscated from traffickers involved in the illegal pet trade or natural history collections. A key goal of species conservation is to reintroduce populations to their historical ranges by relocating individuals under human care into their original habitats. Reintroduction programs play a crucial role in rewilding populations, but over 50% of reintroduction efforts fail (Brichieri-Colombi et al., 2019; Cochran-Biederman et al., 2015; Moseby et al., 2011). The success of these programs often hinges on how well the reintroduced individuals and their associated phenotypes align with the current conditions of the target habitat. Specifically, species-specific traits influence how organisms respond to environmental factors, stressors, and pathogens, thus determining reintroduction success or failure (He et al., 2016; Marr et al., 2018). Ideally, researchers should select individuals under human care who are genetically similar to the original population in the targeted habitat (Pavlova et al., 2017; Kardos et al., 2021). Combining this genetic compatibility with effective introduction programs, such as those outlined by the IUCN (IUCN, 2012), should improve reintroduction outcomes.

Despite the crucial role of reintroduction in species conservation, Ecuadorian zoos lack a system to identify suitable individuals under human care for reintroduction efforts. In the Global South, a significant challenge is the scarcity of resources and expertise for genetic sequencing and analysis (Díaz et al., 2021; Schwartz, 2005). Here, we propose to launch a consortium, the Ecuadorian endangered-species barcoding project (EESBP), that aims to address this issue by establishing a network of collaborations between zoos and natural history museums that will create a genetic library for wild populations.. For example, Quito Zoo annually receives numerous yellow-footed tortoises (*Chelonoidis denticulata*) rescued from illegal trafficking and mascotism (Species360, 2024). A genetic library, incorporating sequences obtained from samples from both natural history museums and field collections, would facilitate the rapid identification of the source population for incoming turtles (e.g., Frandsen et al., 2020; Oklander et al., 2020). Consequently, zoos would be able to reestablish turtle populations where they may be declining. This genetic library would help caregivers reintroduce species by creating species-specific programs that enable individuals of species to acquire essential survival skills (Reading et al., 2013).

Biobanks can support the restoration of ecosystems by technically informing on the origin, identity, and characteristics of species that may be locally extinct or critically endangered (Mooney et al., 2023). Particularly, if a biobank invests in the storage of germ cells, potential breeding programs may be designed to preserve the specific gene pool in the wild (Benirschke, 1984; Bolton et al., 2022; Consortium, 2020). Biobanks are scientific collections of biological samples like tissues, blood, DNA, RNA, among others, and represent valuable genetic resources for a wide range of species. These types of samples are usually cryopreserved, and the associated data is stored in specialized databases (Corrales et al., 2023; Pérez-Espona & Consortium, 2021).

In our work, it has become evident that Ecuador does not have an established biobank for genetic resources of biodiversity, although the National Institute of Biodiversity is currently developing efforts towards establishing one with the support of the Korea International Cooperation Agency (KOICA) and the National Institute of Biological Resources of Korea (NIBR). The joint effort, collaboration, and networking of several institutions working on biodiversity, such as zoos and natural history museums, are fundamental to constituting effective decentralized biobanks of biodiversity. This consortium could provide services and resources for developing breeding manangement and population reintroduction programs, environmental monitoring, among other areas of biotechnological and bioeconomic development (Comizzoli & Wildt, 2017).

### Conclusions

Our study proposed a formula for collaborative networking among zoos, natural history collections, and academia. Through the investigation of olingo genetics, we have demonstrated that inter-institutional collaboration is crucial for rapid species identification for conservation and systematic purposes. Our results revealed the proper taxonomic identification of olingos, which have puzzled zookeepers, veterinarians, and biologists for years. Our analyses strongly support the presence of a single species, *Bassaricyon medius*, in the Quito Zoo population. Importantly, we established a possibly new geographical record in the northern region of Ecuador for *B. m. orinomus,* which could be a potential region for reintroduction of this subspecies. Integrating data from animals under human care and natural history collections allowed us to infer the known diversity of olingos. The possibility of locally sequencing DNA of rare species has profound implications for future studies on species conservation and management, particularly in countries of the Global South. On the one hand, whole genomes can be sequenced to investigate sharp population declines and inbreeding events with precision. On the other hand, accurate taxonomic identification through DNA sequences of such markers as the cytochrome *b* gene can help zookeepers understand key animal behaviors, thus enhancing animal welfare.

## Methods

### Samples and DNA extraction

We collected four whole blood samples from two males (“Munay”, “Macho”) and two females (“Clarissa” and “Lola”) housed in the Quito Zoo. These samples were collected opportunistically during routine veterinary check-ups. The zoo samples were collected from 2021 to 2024 and stored at -20°C at the INABIO collection until genomic DNA extraction. Animals were handled according to the general guidelines of the American Zoological Association (Backues et al., 2011). Sample collection and genetic access were conducted according to Ecuadorian legislation with the corresponding permit (MAATE-DBI-CM-2023-0334). Similarly, we obtained one whole blood sample from the INABIO museum collections, and another sample was collected in 2023 from a male in the Bellavista Cloud Forest Reserve, Ecuador, and deposited in INABIO. We used the Qiagen DNeasy Blood and Tissue kit to extract genomic DNA from these six samples. We evaluated the quality and concentration of DNA with a Qubit 2.0 fluorometer and with an agarose gel electrophoresis method.

### PCR Amplifications and DNA Sequencing

We chose the cytochrome *b* gene (*CYTB*), a widely used molecular marker in mammalian phylogenetics to compare zoo and museum individuals with sequences deposited in GenBank. We used the universal primers MTCB-F (5′-CCHCCATAAATAGGNGAAGG -3′) and MTCB-Rc (5′-WAGAAYTTCAGCTTTGGG-3′) to amplify a 1,146 bp fragment of this gene (Naidu et al., 2012). We followed previous studies for the PCR thermal cycling profile (Naidu et al., 2012). Oxford Nanopore Technologies (ONT) library preparation and sequencing was done at the Nucleic Acid Sequencing Laboratory of the Instituto Nacional de Biodiversidad (INABIO) in Quito, Ecuador. We converted purified products into a high-fidelity sequencing library using the ONT-suggested biochemistry and procedures (reference - even if it’s to a manual or online protocol). Sequencing was performed on a MinION Mk1C with the Rapid Barcoding Kit 96 (SQK-RBK114.96) and Flongle Flow Cells R10.4.1, under the standard protocol. We conducted the demultiplexing step as well as the base-calling process using Dorado v0.9.0., under the super-accurate basecaller (SUP) model. The resulting FASTQ files were filtered at a Q score of 9. Consensus sequences were generated with NGSpecies ID, as described by Sahlin et al. (2021). We deposited the raw fasta sequences in Genbank under the BioProject PRJNA1221930.

### Phylogenetic tree

A total of 17 previously published cytochrome b gene (*CYTB*) sequences were downloaded from NCBI-GenBank, registered with the following accession numbers: AB462085.1 *Procyon lotor*, DQ660299.1 *Bassaricyon alleni*, DQ660300.1 *Bassaricyon medius orinomus*, DQ660301.1 *Bassariscus sumichrasti*, EF107703.1 *Bassaricyon medius orinomus*, EF107704.1 *Bassaricyon medius orinomus*, EF107705.1 Potos flavus, EF107706.1 *Bassaricyon medius medius*, EF107707.1 *Bassaricyon medius medius*, EF107708.1 *Bassaricyon neblina*, EF107709.1 *Bassaricyon neblina*, EF107710.1 *Bassaricyon alleni*, JQ433717.1 Nasua nasua, JX948744.1 Bassaricyon gabbii, MK144297.1 *Bassaricyon medius orinomus*, NC 066722.1 *Bassaricyon neblina*, X94931.1 *Bassaricyon neblina*. We inferred a phylogenetic tree among 23 taxa based on the *CYTB* sequences, after first generating a multiple sequence alignment using MUSCLE (Edgar, 2021). Phylogenetic inference was done in IQ-TREE (Nguyen et al., 2015), using a maximum-likelihood model. We used 1000 ultrafast bootstraps and determined the final phylogeny under the best-fitting substitution model for the alignment using the ModelFinder function (Kalyaanamoorthy et al., 2017). Sequences of *Bassariscus sumichrasti*, *Nasua nasua*, and *Procyon lotor* were used as outgroups to root the phylogenetic tree. The scripts for the sequence alignment and phylogenetic inference are available at https://github.com/dechavezv

To calculate average genomic divergence times across the *Bassaricyon* genus, we used 697 sites of the *CYTB* gene across the 23 analyzed sequences (including *Nasua nasua* and *Procyon lotor* outgroups). We used this alignment with the MCMCTree tool from the PAML 4.8 package (Yang, 2007) and the topology obtained from the phylogenetic analysis to estimate divergence times. The HKY + G model (Hasegawa et al., 1985) was selected by ModelFinder in IQ-Tree to estimate the divergence times. We ran the MCMC for 2,200,000 iterations, sampling every 2nd iteration, and discarding the first 200,000 iterations as burn-in. We used fossil priors to calibrate specific nodes of the phylogeny (Table S2).

## Supplementary Information

**Figure S1.**
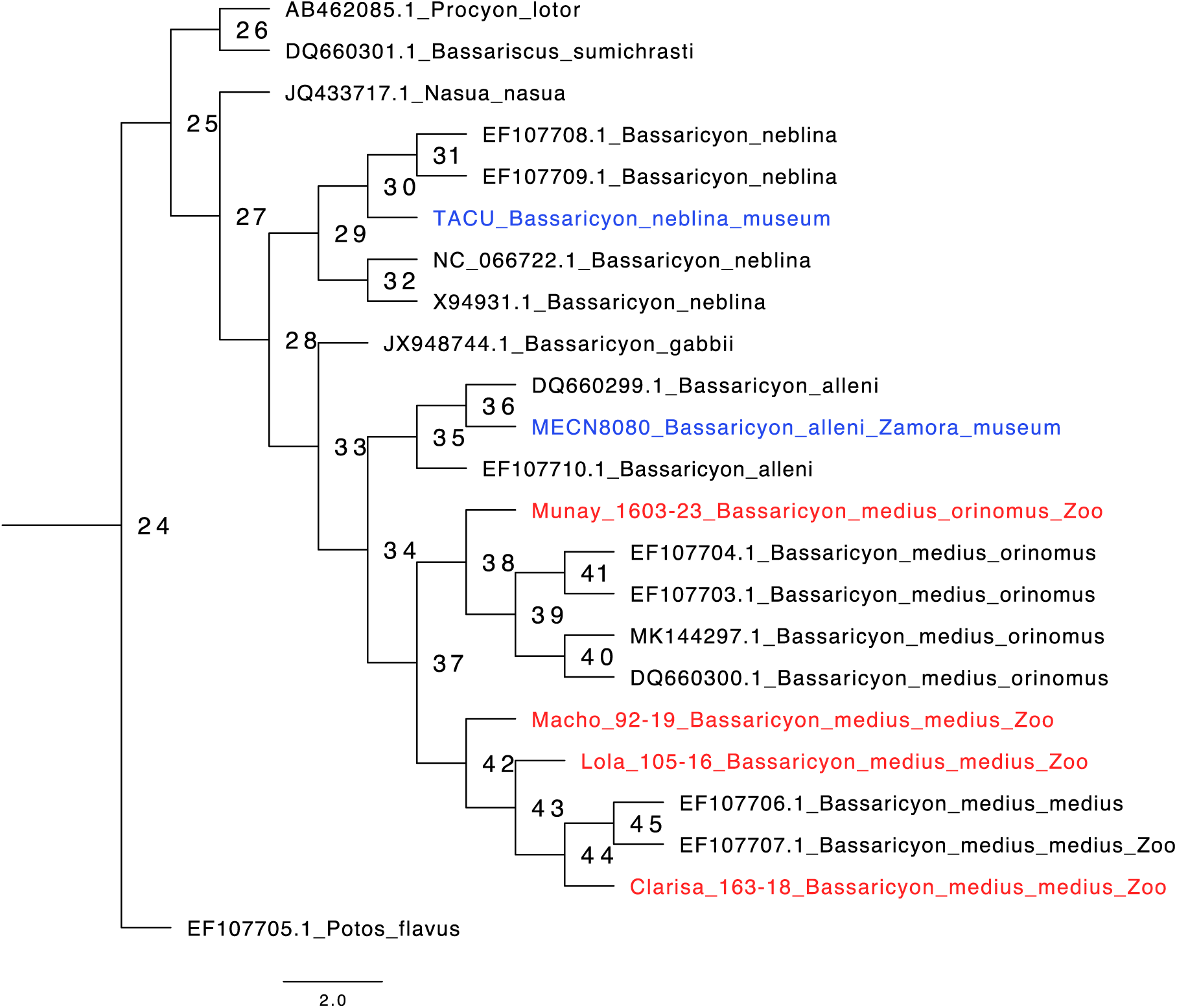
Maximum likelihood tree inferred using IQ-TREE which includes the studied samples of olingos. The number on the nodes corresponds to the numbers shown in TableS1. Calibration priors of credible intervals are in Table S2. Blue-labeled branches are samples from the INABIO. Red-labeled branches are sequences from individuals at the Quito Zoo.

**Table S1.** Mean divergence times and 95% highest posterior densities estimated using MCMCTree at the nodes numbered in Figure S1 and corresponding to the Figure 1.

| Node in the Tree | Mean Divergence Time (95% highest posterior density) |
| --- | --- |
| N24 | 16.1967 (13.6809, 18.6802) |
| N25 | 13.6808 (11.1741, 16.8552) |
| N26 | 11.4999 (11.0013, 12.0014) |
| N27 | 7.199 (5.5062, 8.9066) |
| N28 | 5.6384 (3.2537, 8.1383) |
| N29 | 3.0073 (0.8309, 6.0426) |
| N30 | 1.6286 (0.0591, 3.3974) |
| N31 | 0.7416 (0.0000, 1.9532) |
| N32 | 1.1868 (0.0000, 2.6744) |
| N33 | 4.022 (2.5247, 6.1835) |
| N34 | 2.6502 (2.4998, 2.7998) |
| N35 | 1.6606 (0.4460, 2.7106) |
| N36 | 0.7824 (0.0000, 1.9487) |
| N37 | 2.33 (1.7441, 2.7810) |
| N38 | 1.7949 (0.8708, 2.6082) |
| N39 | 1.2973 (0.3500, 2.2402) |
| N40 | 0.6175 (0.0000, 1.5188) |
| N41 | 0.6191 (0.0000, 1.5257) |
| N42 | 1.7953 (0.8698, 2.6050) |
| N43 | 1.2978 (0.3501, 2.2379) |
| N44 | 0.8346 (0.0574, 1.7170) |
| N45 | 0.4029 (0.0000, 1.1151) |

**Table S2.** Set of fossil calibration priors (minimum and maximum node ages) used for the MCMCTree analyses.

| Node<br>on<br>tree | Node | Node<br>minimum<br>age (Ma) | Node<br>maximum<br>age (Ma) | Evidence | Prior model |
| --- | --- | --- | --- | --- | --- |
| 1 | Potos flavus split | 13.7 | 18.7 | (Faurby et al., 2024) | Lognormal |
| 2 | TMRCAs *Procyon* +<br>*Bassaricyus* | 11 | 12 | (Faurby et al., 2024; Koepfli et al., 2007) | Lognormal |
| 3 | *Nasua nasua* split | 5.5 | 8.9 | (Faurby et al., 2024) | Lognormal |
| 4 | *Bassaricyon alleni* - split<br>*Bassaricyon medius* | 2.5 | 2.8 | (Faurby et al., 2024; Koepfli et al., 2007) | Lognormal |

Note – Average genomic divergence time estimates are shown in Figure 1 and Table S1.

## Acknowledgements

We thank Santiago Ron and Omar Torres from the Pontificia Universidad Católica del Ecuador (PUCE) for providing the laboratory space and equipmentto conduct the DNA extraction of the olingos analyzed in this study. We also thank Nicole Omasa and Betsabé Trujillo from the Fundación Zoológica del Ecuador for helping to collect the sample from the olingos sequenced in this study. We thank Carolina Saenz from Tueri at San Francisco University for providing information about the olingo named “Macho”. We thank Santiago Cárdenas from the Amaru Bioparque for sharing information on the olingo named “Clarissa”. We thank Jorge Brito from INABIO for including the olingo sample in the mammalian collection. We thank Klaus-Peter Koepfli for proofreading the manuscript. We acknowledge the support from the Dirección de Investigación at PUCE with the grant PEP QINV0502-IINV529010100 to D.E.C.

## Author Contributions

D.E.C., J.C.C, D.M., M.B.C. M.B, C.M.P, P.J.V designed the study and performed the analyses. D.E.C. C. M.P and J.C.C processed the sequencing reads and performed the read base-calling, phylogenetic analysis divergence time analysis. J.C.C, M.B.C, D.R.-B, P.L, D.M organized and coordinated the sample collection and sequencing of the olingos. The manuscript was written by D.E.C., C.M.P and P.J.V. with subsequent contributions and the final approval by all authors.

## References

Alves, R. J. V., Weksler, M., Oliveira, J. A., Buckup, P. A., Pombal, J. P., Santana, H. R. G.,…Caramaschi, U. (2018). Brazilian legislation on genetic heritage harms Biodiversity Convention goals and threatens basic biology research and education. An Acad Bras Cienc, 90(2), 1279–1284. 10.1590/0001-3765201820180460

Andrews, G., Fan, K., Pratt, H. E., Phalke, N., Karlsson, E. K., Lindblad-Toh, K.,… Consortium§, Z. (2023). Mammalian evolution of human cis-regulatory elements and transcription factor binding sites. Science, 380(6643), eabn7930. 10.1126/science.abn7930

Backues, K., Clyde, V., Denver, M., Fiorello, C., Hilsenroth, R., Lamberski, N.,… Veterinarians, E. C. A. A. o. Z. (2011). Guidelines for zoo and aquarium veterinary medical programs and veterinary hospitals. J Zoo Wildl Med, 42(1), 176–192.

Benirschke, K. (1984). The frozen zoo concept. Zoo Biology, 3(4), 325–328.

Berta, A. (1987). Origin, diversification, and zoogeography of the South American Canidae. Fieldiana. Zoology[FIELDIANA ZOOL.].

Bolaños-Villegas, P., Cabrerizo, F. M., Brown, F. D., Zancan, P., Barrera, J. F., González-Muñoz, P. A.,…Vargas-Balda, R. (2020). Latin America: reduced S&T investment puts sustainable development at risk. ScienceOpen Preprints.

Bolton, R. L., Mooney, A., Pettit, M. T., Bolton, A. E., Morgan, L., Drake, G. J.,…Hvilsom, C. (2022). Resurrecting biodiversity: advanced assisted reproductive technologies and biobanking. Reprod Fertil, 3(3), R121–R146. 10.1530/RAF-22-0005

Brichieri-Colombi, T. A., Lloyd, N. A., McPherson, J. M., & Moehrenschlager, A. (2019). Limited contributions of released animals from zoos to North American conservation translocations. Conserv Biol, 33(1), 33–39. 10.1111/cobi.13160

Brito, J., García, R., Castellanos, F. X., Gavilanes, G., Curay, J., Carrión-Olmedo, J. C.,…Pinto, C. M. (2024). Two new species of Thomasomys (Cricetidae: Sigmodontinae) from the western Andes of Ecuador and an updated phylogenetic hypothesis for the genus. Vertebrate Zoology, 74, 709–734.

Bush, E. R., Baker, S. E., & Macdonald, D. W. (2014). Global trade in exotic pets 2006–2012. Conservation Biology, 28(3), 663–676.

Chacón-Pacheco, J., Bassa-Hernández, D. J., & Ramírez-Chaves, H. E. (2019). New record of Bassaricyon medius in the Colombian Caribbean. Therya, 10(2), 201–205.

Chavez, D. E., Gronau, I., Hains, T., Dikow, R. B., Frandsen, P. B., Figueiro, H. V.,…Wayne, R. K. (2022). Comparative genomics uncovers the evolutionary history, demography, and molecular adaptations of South American canids. Proceedings of the National Academy of Sciences of the United States of America, 119(34), Article e2205986119. 10.1073/pnas.2205986119

Che-Castaldo, J. P., Grow, S. A., & Faust, L. J. (2018). Evaluating the Contribution of North American Zoos and Aquariums to Endangered Species Recovery. Sci Rep, 8(1), 9789. 10.1038/s41598-018-27806-2

Christmas, M. J., Kaplow, I. M., Genereux, D. P., Dong, M. X., Hughes, G. M., Li, X.,… Consortium§, Z. (2023). Evolutionary constraint and innovation across hundreds of placental mammals. Science, 380(6643), eabn3943. 10.1126/science.abn3943

Coates, A. G., & Obando, J. A. (1996). The geological evolution of the Central American isthmus. In: Jackson, J.B.C., Budd, A.F., Coates, A.G. (Eds.), Evolution and Evironment in Tropical America. In: University of Chicago Press, Chicago, IL, pp. 21–56.

Cochran-Biederman, J. L., Wyman, K. E., French, W. E., & Loppnow, G. L. (2015). Identifying correlates of success and failure of native freshwater fish reintroductions. Conserv Biol, 29(1), 175–186. 10.1111/cobi.12374

Comizzoli, P., & Wildt, D. E. (2017). Cryobanking biomaterials from wild animal species to conserve genes and biodiversity: relevance to human biobanking and biomedical research. Biobanking of human biospecimens: Principles and practice, 217-235.

Consortium, Z. (2020). A comparative genomics multitool for scientific discovery and conservation. Nature, 587(7833), 240–245. 10.1038/s41586-020-2876-6

Corrales, C., Luciano, S., & Astrin, J. J. (2023). Biodiversity biobanks: A landscape analysis. ARPHA Preprints, 4, e103105.

Dados, N., & Connell, R. (2012). The global south. Contexts, 11(1), 12–13.

Dumancas, G. G., Smith, K. F. K., Bugayong-Janagap, A. M., Zamora, P. R. F. C., Ferriols, V. M. E. N., Liwag, A. G.,…Rodney Jr, T. (2023). Challenges and opportunities in establishing a regional biobank center in a developing country: A case from the Philippines. Health Policy and Technology, 100822.

Díaz, M., Jarrín-V, P., Simarro, R., Castillejo, P., Tenea, G. N., & Molina, C. A. (2021). The Ecuadorian Microbiome Project: a plea to strengthen microbial genomic research. Neotropical Biodiversity, 7(1), 223–237.

Edgar, R. C. (2021). MUSCLE v5 enables improved estimates of phylogenetic tree confidence by ensemble bootstrapping. BioRxiv, 2021.2006. 2020.449169.

Eizirik, E., Murphy, W. J., Koepfli, K. P., Johnson, W. E., Dragoo, J. W., Wayne, R. K., & O’Brien, S. J. (2010). Pattern and timing of diversification of the mammalian order Carnivora inferred from multiple nuclear gene sequences. Molecular Phylogenetics and Evolution, 56(1), 49–63. 10.1016/j.ympev.2010.01.033

Ekblom, R., Brechlin, B., Persson, J., Smeds, L., Johansson, M., Magnusson, J.,…Ellegren, H. (2018). Genome sequencing and conservation genomics in the Scandinavian wolverine population. Conserv Biol, 32(6), 1301–1312. 10.1111/cobi.13157

Faurby, S., Silvestro, D., Werdelin, L., & Antonelli, A. (2024). Reliable biogeography requires fossils: insights from a new species-level phylogeny of extinct and living carnivores. Proc Biol Sci, 291(2028), 20240473. 10.1098/rspb.2024.0473

Foley, N. M., Mason, V. C., Harris, A. J., Bredemeyer, K. R., Damas, J., Lewin, H. A.,… Consortium‡, Z. (2023). A genomic timescale for placental mammal evolution. Science, 380(6643), eabl8189. 10.1126/science.abl8189

Frandsen, P., Fontsere, C., Nielsen, S. V., Hanghøj, K., Castejon-Fernandez, N., Lizano, E.,…Hvilsom, C. (2020). Targeted conservation genetics of the endangered chimpanzee. Heredity (Edinb), 125(1-2), 15–27. 10.1038/s41437-020-0313-0

GBIF.org. GBIF Occurrence Download. Retrieved January 31 from

GBIF.org. (2025). GBIF Occurrence Download 10.15468/dl.fv5dfz. In.

Harrington, L. A. (2015). International commercial trade in live carnivores and primates 2006–2012: response to Bush et al. 2014. Conservation Biology, 29(1), 293-296.

Hasegawa, M., Kishino, H., & Yano, T. A. (1985). DATING OF THE HUMAN APE SPLITTING BY A MOLECULAR CLOCK OF MITOCHONDRIAL-DNA. Journal of Molecular Evolution, 22(2), 160–174. 10.1007/bf02101694

He, X., Johansson, M. L., & Heath, D. D. (2016). Role of genomics and transcriptomics in selection of reintroduction source populations. Conserv Biol, 30(5), 1010–1018. 10.1111/cobi.12674

Helgen, K. K., R.Pinto, C. Schipper, J. (2016). Bassaricyon medius in IUCN 2017. The IUCN Red List of Threatened Species. Version 2017.3 www.iucnredlist.org. Accessed on January 27 of 2025. In.

Helgen, K. M., Pinto, C. M., Kays, R., Helgen, L. E., Tsuchiya, M. T. N., Quinn, A.,…Maldonado, J. E. (2013). Taxonomic revision of the olingos (*Bassaricyon*), with description of a new species, the Olinguito. Zookeys(324), 1-83. 10.3897/zookeys.324.5827

Iucn, R. (2012). IUCN guidelines for reintroductions and other conservation translocations. IUCN: Gland, Switzerland.

Jarrín-V, P., Falconí, F., Cango, P., & Ramos-Martin, J. (2021). Knowledge gaps in Latin America and the Caribbean and economic development. World development, 146, 105602.

Jayathissa, P., & Rupasinghe, A. (2024). Exploring DNA Analysis Methods and Genetic Research Applications in Low and Middle-Income Nations: A Study of Sri Lanka. Available at SSRN 4784002.

Kalyaanamoorthy, S., Minh, B. Q., Wong, T. K. F., von Haeseler, A., & Jermiin, L. S. (2017). ModelFinder: fast model selection for accurate phylogenetic estimates. Nature Methods, 14(6), 587-+. 10.1038/nmeth.4285

Khan, S., Nabi, G., Ullah, M. W., Yousaf, M., Manan, S., Siddique, R., & Hou, H. (2016). Overview on the Role of Advance Genomics in Conservation Biology of Endangered Species. Int J Genomics, 2016, 3460416. 10.1155/2016/3460416

Koepfli, K. P., Gompper, M. E., Eizirik, E., Ho, C. C., Linden, L., Maldonado, J. E., & Wayne, R. K. (2007). Phylogeny of the Procyonidae (Mammalia: Carnivora): Molecules, morphology and the Great American Interchange. Molecular Phylogenetics and Evolution, 43(3), 1076–1095. 10.1016/j.ympev.2006.10.003

Kyriazis, C. C., Robinson, J. A., & Lohmueller, K. E. (2023). Using Computational Simulations to Model Deleterious Variation and Genetic Load in Natural Populations. Am Nat, 202(6), 737–752. 10.1086/726736

Lawson, C., Rourke, M., & Humphries, F. (2022). Access and Benefit Sharing of Genetic Resources, Information, and Traditional Knowledge. Routledge.

Marr, M. M., Brace, S., Schreve, D. C., & Barnes, I. (2018). Identifying source populations for the reintroduction of the Eurasian beaver, Castor fiber L. 1758, into Britain: evidence from ancient DNA. Sci Rep, 8(1), 2708. 10.1038/s41598-018-21173-8

Miranda, A., Altamirano, A., Cayuela, L., Pincheira, F., & Lara, A. (2015). Different times, same story: Native forest loss and landscape homogenization in three physiographical areas of south-central of Chile. Applied Geography, 60, 20–28. 10.1016/j.apgeog.2015.02.016

Miranda, R., Escribano, N., Casas, M., Pino-del-Carpio, A., & Villarroya, A. (2023). The role of zoos and aquariums in a changing world. Annual Review of Animal Biosciences, 11(1), 287–306.

Mooney, A., Ryder, O. A., Houck, M. L., Staerk, J., Conde, D. A., & Buckley, Y. M. (2023). Maximizing the potential for living cell banks to contribute to global conservation priorities. Zoo Biol, 42(6), 697–708. 10.1002/zoo.21787

Moseby, K., Read, J., Paton, D., Copley, P., Hill, B., & Crisp, H. (2011). Predation determines the outcome of 10 reintroduction attempts in arid South Australia. Biological Conservation, 144(12), 2863–2872.

Naidu, A., Fitak, R. R., Munguia-Vega, A., & Culver, M. (2012). Novel primers for complete mitochondrial cytochrome b gene sequencing in mammals. Mol Ecol Resour, 12(2), 191–196. 10.1111/j.1755-0998.2011.03078.x

Nguyen, L. T., Schmidt, H. A., von Haeseler, A., & Minh, B. Q. (2015). IQ-TREE: A Fast and Effective Stochastic Algorithm for Estimating Maximum-Likelihood Phylogenies. Molecular Biology and Evolution, 32(1), 268–274. 10.1093/molbev/msu300

O’Dea, A., Lessios, H. A., Coates, A. G., Eytan, R. I., Restrepo-Moreno, S. A., Cione, A. L.,…Jackson, J. B. C. (2016). Formation of the Isthmus of Panama. Science Advances, 2(8), Article e1600883. 10.1126/sciadv.1600883

Oklander, L. I., Caputo, M., Solari, A., & Corach, D. (2020). Genetic assignment of illegally trafficked neotropical primates and implications for reintroduction programs. Sci Rep, 10(1), 3676. 10.1038/s41598-020-60569-3

Ortega, J. A., Cisneros-Heredia, D. F., Camper, J. D., Romero-Carvajal, A., Negrete, L., & Ron, S. R. (2025). Systematics of minute strabomantid frogs allocated to the genus Noblella (Amphibia: Anura) with description of a new genus, seven new species, and insights into historical biogeography. Zoological Journal of the Linnean Society, 203(1), zlae162.

Patrick, P. G., & Tunnicliffe, S. D. (2013). Rationale for the Existence of Zoos. Zoo Talk, 19-35.

Pomerantz, A., Peñafiel, N., Arteaga, A., Bustamante, L., Pichardo, F., Coloma, L. A.,…Prost, S. (2018). Real-time DNA barcoding in a rainforest using nanopore sequencing: opportunities for rapid biodiversity assessments and local capacity building. GigaScience, 7(4), giy033.

Poo, S., Whitfield, S. M., Shepack, A., Watkins-Colwell, G. J., Nelson, G., Goodwin, J.,…Chakrabarty, P. (2022). Bridging the Research Gap between Live Collections in Zoos and Preserved Collections in Natural History Museums. Bioscience, 72(5), 449–460. 10.1093/biosci/biac022

Powell, D. M., Meyer, T. G., & Duncan, M. (2023). By Bits and Pieces: The Contributions of Zoos and Aquariums to Science and Society via Biomaterials. Journal of zoological and botanical gardens, 4(1), 277–287. 10.3390/jzbg4010023

Pozo, G., Albuja-Quintana, M., Larreátegui, L., Gutiérrez, B., Fuentes, N., Alfonso-Cortés, F., & Torres, M. L. (2024). First whole-genome sequence and assembly of the Ecuadorian brown-headed spider monkey (Ateles fusciceps fusciceps), a critically endangered species, using Oxford Nanopore Technologies. G3 (Bethesda), 14(3). 10.1093/g3journal/jkae014

Pérez-Espona, S., & Consortium, C. (2021). Conservation-focused biobanks: a valuable resource for wildlife DNA forensics. Forensic Science International: Animals and Environments, 1, 100017.

Ramírez-Chaves, H. E., Ossa-López, P. A., Velásquez-Guarín, D., Colmenares-Pinzón, J., Noguera-Urbano, E. A., Mejía-Fontecha, I. Y.,…Suárez-Castro, A. F. (2022). New genetic information and geographic distribution of charismatic carnivores: the olingos (Procyonidae: Bassaricyon) in Colombia. Mammalian Biology, 102(5), 2045–2059.

Robinson, J., Kyriazis, C. C., Yuan, S. C., & Lohmueller, K. E. (2023). Deleterious Variation in Natural Populations and Implications for Conservation Genetics. Annual Review of Animal Biosciences, 11, 93–114. 10.1146/annurev-animal-080522-093311

Robinson, J. A. K., C.C, Nigenda-Morales, S. F., Beichman, A. C., Rojas-Bracho, L., Robertson, K. M., Fontaine, M. C., … Morin, P. A. (2022). The critically endangered vaquita is not doomed to extinction by inbreeding depression. 376(6593), 635–639.

Rubinoff, D., Reil, J. B., Osborne, K. H., Gregory, C. J., Geib, S. M., & Dupuis, J. R. (2020). Phylogenomics reveals conservation challenges and opportunities for cryptic endangered species in a rapidly disappearing desert ecosystem. Biodiversity and Conservation, 29, 2185–2200.

Saavedra-Rodríguez, C. A., & Velandia-Perilla, J. H. (2011). Bassaricyon gabbii Allen, 1876 (Carnivora: Procyonidae): New distribution point on western range of Colombian Andes. Check List, 7(4), 505-507.

Schwartz, M. K. (2005). Guidelines on the use of molecular genetics in reintroduction programs. In: The EU LIFE-Nature Projects to guidelines for the reintroduction of threatened species; Caramanico Terme, Italy; March 21-22, 2005. p. 51-58, 51-58.

Shaw, R. E., Brockett, B., Pierson, J. C., Sarre, S. D., Doyle, P., Cliff, H. B.,…Parrott, M. L. (2024). Building meaningful collaboration in conservation genetics and genomics. Conservation Genetics, 1-19.

Species360. (2024). Species360 Zoological Information Management System (ZIMS). In: zims.Species360.org.

Sullivan, J., Arellano, E., & Rogers, D. S. (2000). Comparative phylogeography of mesoamerican highland rodents: Concerted versus independent response to past climatic fluctuations. American Naturalist, 155(6), 755–768. 10.1086/303362

Sullivan, P. F., Meadows, J. R. S., Gazal, S., Phan, B. N., Li, X., Genereux, D. P.,… Consortium§, Z. (2023). Leveraging base-pair mammalian constraint to understand genetic variation and human disease. Science, 380(6643), eabn2937. 10.1126/science.abn2937

Vilaça, S. T., Vidal, A. F., Pavan, A. C. D. O., Silva, B. M., Carvalho, C. S., Povill, C.,…Mendes, I. S. (2024). Leveraging genomes to support conservation and bioeconomy policies in a megadiverse country. Cell Genomics, 4(11).

Wang, Y., Zhao, Y., Bollas, A., & Au, K. F. (2021). Nanopore sequencing technology, bioinformatics and applications. Nat Biotechnol, 39(11), 1348–1365. 10.1038/s41587-021-01108-x

Webb, D. S. (1991). Ecography and the Great American Interchange. 3, 226–280.

Webb, S. D. (2006). The Great American Biotic Interchange: Patterns and processes. Annals of the Missouri Botanical Garden, 93(2), 245–257. <Go to ISI>://WOS:000240071400005

Webb, S. D., & Rancy, A. (1996). Late Cenozoic evolution of the Neotropical mammal fauna. In J. B. C. Jackson, A. F. Budd, & A. G. Coates (Eds.), Evolution and Environment in Tropical America (pp. 335–358). The University of Chicago Press, Chicago.

Webb, S. D. C. F. p. d. (1978). A History of Savanna Vertebrates in the New World. Part II: South America and the Great Interchange. Annual Review of Ecology and Systematics, 9, 393–426.

Weidener, L., & Spreckelsen, C. (2024). Decentralized science (DeSci): definition, shared values, and guiding principles. Frontiers in Blockchain, 7, 1375763.

Wilder, A. P., Supple, M. A., Subramanian, A., Mudide, A., Swofford, R., Serres-Armero, A.,… Consortium‡, Z. (2023). The contribution of historical processes to contemporary extinction risk in placental mammals. Science, 380(6643), eabn5856. 10.1126/science.abn5856

Yang, Z. H. (2007). PAML 4: Phylogenetic analysis by maximum likelihood. Molecular Biology and Evolution, 24(8), 1586–1591. 10.1093/molbev/msm088

